# When Geometric Deep Learning Meets Pretrained Protein Language Models

**DOI:** 10.1101/2023.01.05.522958

**Authors:** Fang Wu, Yu Tao, Dragomir Radev, Jinbo Xu

## Abstract

Geometric deep learning has recently achieved great success in non-Euclidean domains, and learning on 3D structures of large biomolecules is emerging as a distinct research area. However, its efficacy is largely constrained due to the limited quantity of structural data. Meanwhile, protein language models trained on substantial 1D sequences have shown burgeoning capabilities with scale in a broad range of applications. Nevertheless, no preceding studies consider combining these different protein modalities to promote the representation power of geometric neural networks. To address this gap, we make the foremost step to integrate the knowledge learned by well-trained protein language models into several state-of-the-art geometric networks. Experiments are evaluated on a variety of protein representation learning benchmarks, including protein-protein interface prediction, model quality assessment, protein-protein rigid-body docking, and binding affinity prediction, leading to an overall improvement of 20% over baselines and the new state-of-the-art performance. Strong evidence indicates that the incorporation of protein language models’ knowledge enhances geometric networks’ capacity by a significant margin and can be generalized to complex tasks.

## 1 Introduction

Macromolecules (*e*.*g*., proteins, RNAs, or DNAs) are essential to biophysical processes. While they can be represented using lower-dimensional representations such as linear sequences (1D) or chemical bond graphs (2D), a more intrinsic and informative form is the three-dimensional geometry [97]. 3D shapes are critical to not only understanding the physical mechanisms of action but also answering a number of questions associated with drug discovery and molecular design [86]. As a consequence, tremendous efforts in structural biology have been devoted to deriving insights from their conformations [53, 55, 96].

With the rapid advances of deep learning (DL) techniques, it has been an attractive challenge to represent and reason about macromolecules’ structures in the 3D space. In particular, different sorts of 3D information including bond lengths and dihedral angles play an essential role. In order to encode them, a number of 3D geometric graph neural networks (GGNNs) or CNNs [9, 42, 44, 45, 81, 94, 95] have been proposed, and simultaneously achieve several crucial properties of Euclidean geometry such as E(3) or SE(3) equivariance and symmetry. Notably, they are important constituents of geometric deep learning (GDL), an umbrella term that generalizes networks to Euclidean or non-Euclidean domains [5].

Meanwhile, the anticipated growth of sequencing promises unprecedented data on natural sequence diversity. The abundance of 1D amino acid sequences has spurred increasing interest in developing protein language models at the scale of evolution, such as the series of ESM [54, 60, 71] and ProtTrans [25]. These protein language models are capable of capturing information about secondary and tertiary structures and can be generalized across a broad range of downstream applications. To be explicit, they have recently been demonstrated with strong capabilities in uncovering protein structures [54], predicting the effect of sequence variation on function [60], learning inverse folding [40] and many other general purposes [71].

Despite the fruitful progress in protein language models, no prior studies have considered enhancing GGNNs’ ability by leveraging the knowledge of those protein language models. This is nontrivial because compared to sequence learning, 3D structures are much harder to obtain and thus less prevalent. Consequently, learning on the structure of proteins leads to a reduced amount of training data. For example, the SAbDab database [23] merely has 3K antibody-antigen structures without duplicate. The SCOPe database [16] has 226K annotated structures, and the SIFTS database [89] comprises around 220K annotated enzyme structures. These numbers are orders of magnitude lower than the data set sizes that can inspire major breakthroughs in the deep learning community. In contrast, while the Protein Data Bank (PDB) [12] possesses approximately 182K macromolecule structures, databases like Pfam [61] and UniParc [8] contains more than 47M and 250M protein sequences respectively.

In addition to the data size, the benefit of protein sequence to structure learning also has solid evidence and theoretical support. Remarkably, the idea that biological function and structures are documented in the statistics of protein sequences selected through evolution has a long history [2, 98]. The unobserved variables that decide a protein’s fitness, including structure, function, and stability, leave a record in the distribution of observed natural sequences [35]. Those protein language models use self-supervision to unlock the information encoded in protein sequence variations, which is also beneficial for GGNNs.

Accordingly, in this paper, we propose to promote the capacity of GGNNs with the knowledge learned by protein language models (see Figure 1). The improvements come from two major lines. Firstly, GGNNs can benefit from the information that emerges in the learned representations of those protein language models on fundamental properties of proteins, including secondary structures, contacts, and biological activity. This kind of knowledge may be difficult for GGNNs to be aware of and learn in a specific downstream task. To confirm this claim, we conduct a toy experiment to demonstrate that conventional graph connectivity mechanisms prevent existing GGNNs from being cognizant of residues’ absolute and relative positions in the protein sequence. Secondly and more intuitively, protein language models serve as an alternative way of enriching GGNNs’ training data and allow GGNNs to be exposed to more different families of proteins, which thereby greatly strengthens GGNNs’ generalization capability.

**Fig. 1:**
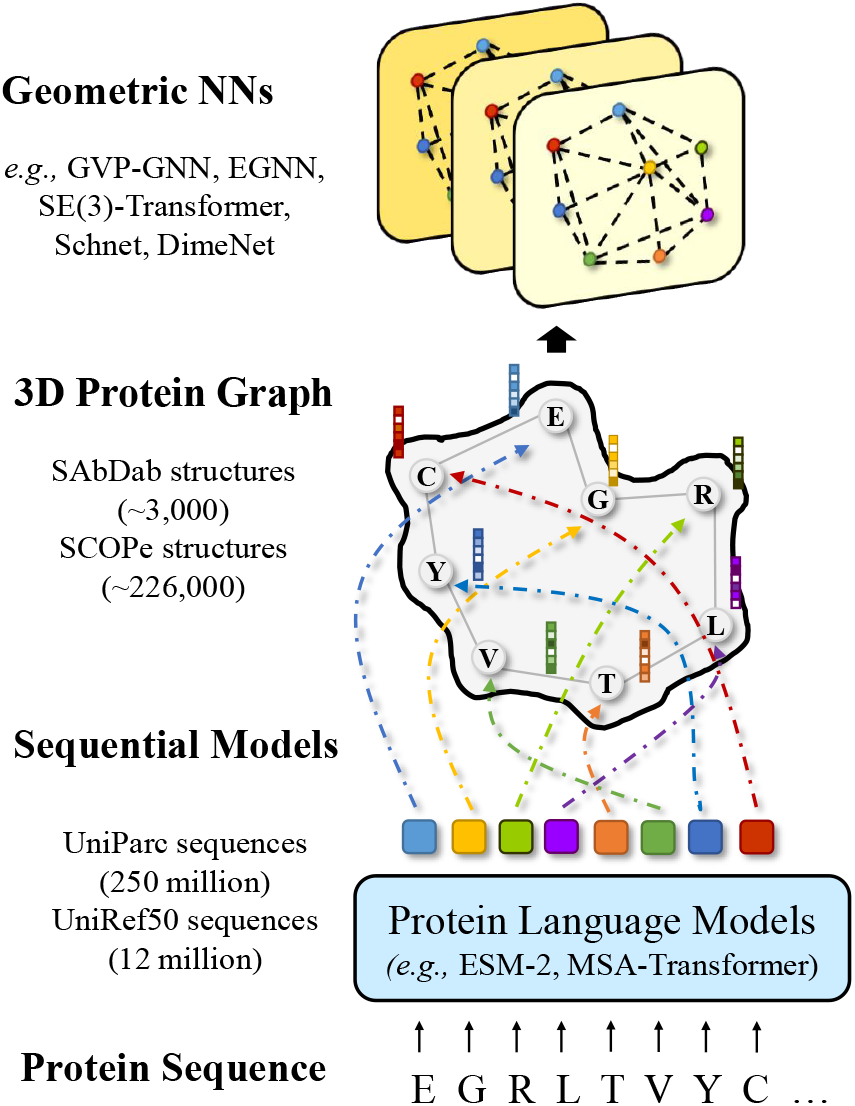
Illustration of our proposed framework to strengthen GGNNs with knowledge of protein language models. The protein sequence is first forwarded into a pretrained protein language model to extract per-residue representations, which are then used as the node feature in 3D protein graphs for GGNNs.

We examine our hypothesis across a wide range of benchmarks, containing model quality assessment, protein-protein interface prediction, protein-protein rigid-body docking, and ligand binding affinity prediction. Extensive experiments show that the incorporation and combination of pretrained protein language models’ knowledge significantly improve GGNNs’ performance for various problems, which require distinct domain knowledge. By utilizing the unprecedented view into the language of protein sequences provided by powerful protein language models, GGNNs promise to augment our understanding of a vast database of poorly understood protein structures. Our work provides a new perspective on protein representation learning and hopes to shed light on how to bridge the gap between the thriving geometric deep learning and mature protein language models, which independently leverage different modalities of proteins.

## 2 Experiments

Our toy experiments illustrate that existing GGNNs are unaware of the positional order inside the protein sequences. Taking a step further, we show in this section that incorporating knowledge learned by large-scale protein language models can robustly enhance GGNN’s capacity in a wide variety of downstream tasks.

### 2.1 Tasks and Datasets

- **Model Quality Assessment** (MQA) aims to select the best structural model of a protein from a large pool of candidate structures and is an essential step in structure prediction [17]. For a number of recently solved but unreleased structures, structure generation programs produce a large number of candidate structures. MQA approaches are evaluated by their capability of predicting the GDT-TS score of a candidate structure compared to the experimentally solved structure of that target. Its database is composed of all structural models submitted to the Critical Assessment of Structure Prediction (CASP) [52] over the last 18 years. The data is split temporally by competition year. MQA is similar to the Protein Structure Ranking (PSR) task introduced by Townshend et al. [86].
- **Protein-protein Rigid-body Docking** (PPRD) computationally predicts the 3D structure of a protein-protein complex from the individual unbound structures. It assumes that no conformation change within the proteins happens during binding. We leverage Docking Benchmark 5.5 (DB5.5) [91] as the database. It is a gold standard dataset in terms of data quality and contains 253 structures.
- **Protein-protein Interface** (PPI) investigates whether two amino acids will contact when their respective proteins bind. It is an important problem in understanding how proteins interact with each other, *e*.*g*., antibody proteins recognize diseases by binding to antigens. We use the Database of Interacting Protein Structures (DIPS), a comprehensive dataset of protein complexes mined from the PDB [85], and randomly select 15K samples for evaluation.
- **Ligand Binding Affinity** (LBA) is an essential task for drug discovery applications. It predicts the strength of a candidate drug molecule’s interaction with a target protein. Specifically, we aim to forecast *pK* = − log_10_ *K*, where *K* is the binding affinity in Molar units. We use the PDBbind database [57, 93], a curated database containing protein-ligand complexes from the PDB and their corresponding binding strengths. The protein-ligand complexes are split such that no protein in the test dataset has more than 30% or 60% sequence identity with any protein in the training dataset.

### 2.2 Experimental Setup

We evaluate our proposed framework on the instances of several state-of-the-art geometric networks, using Pytorch [66] and PyG [28] on four standard protein benchmarks. For MQA, PPI, and LBA, we use backbones that have been carefully described in Section 5, *i*.*e*., GVP-GNN, EGNN, and Molformer. For PPRD, we utilize the state-of-the-art deep learning model, EquiDock [32], as the backbone. It approximates the binding pockets and obtains the docking poses using keypoint matching and alignment. For more experimental details, please refer to Appendix A.

### 2.3 Results

Results are reported with mean ± standard deviation over three repeated runs, where the best performance is in bold. The column of ‘PLM’ indicates whether the protein language model is used.

#### 2.3.1 Single-protein Representation Task

For MQA, we document Spearman correlation (*R*_*S*_), Pearson’s correlation (*R*_*P*_), and Kendall rank correlation (*K*_*R*_) (see Table 1). The introduction of protein language models has brought a significant average increase of 32.63% and 55.71% in global and mean *R*_*S*_, of 34.66% and 58.75% in global and mean *R*_*P*_, and of 43.21% and 63.20% in global and mean *K*_*R*_ respectively. With the aid of language models, GVP-GNN achieves the optimal global *R*_*S*_, global *R*_*P*_, and *K*_*R*_ of 84.92%, 85.44%, and 67.98% separately.

**Table 1:**
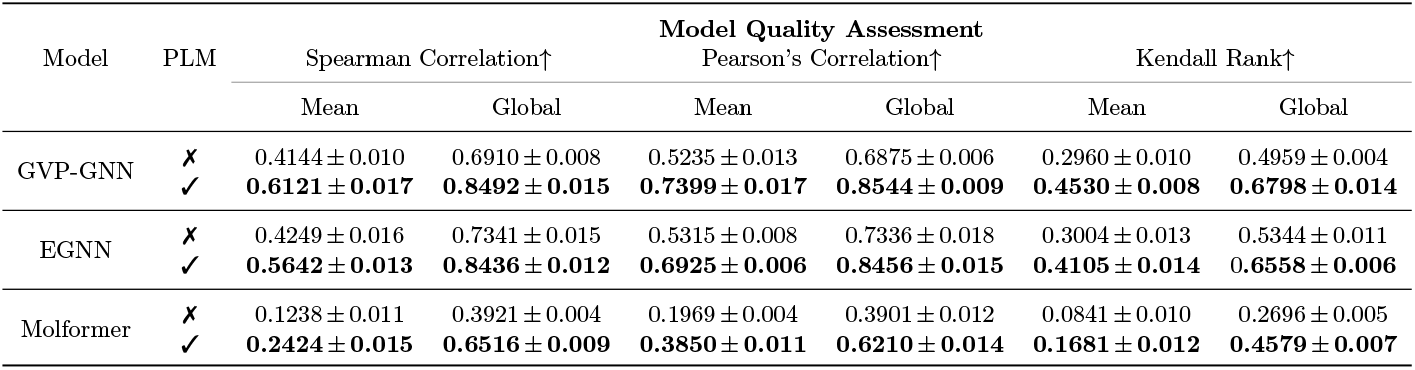
Results on MQA.

| Model | PLM | Model Quality Assessment |  |  |  |  |  |
| --- | --- | --- | --- | --- | --- | --- | --- |
| | | Spearman Correlation↑ | | Pearson's Correlation↑ | | Kendall Rank↑ | |
|  |  | Mean | Global | Mean | Global | Mean | Global |
| GVP-GNN | × | 0.4144 ± 0.010 | 0.6910 ± 0.008 | 0.5235 ± 0.013 | 0.6875 ± 0.006 | 0.2960 ± 0.010 | 0.4959 ± 0.004 |
| | ✓ | **0.6121 ± 0.017** | **0.8492 ± 0.015** | **0.7399 ± 0.017** | **0.8544 ± 0.009** | **0.4530 ± 0.008** | **0.6798 ± 0.014** |
| EGNN | × | 0.4249 ± 0.016 | 0.7341 ± 0.015 | 0.5315 ± 0.008 | 0.7336 ± 0.018 | 0.3004 ± 0.013 | 0.5344 ± 0.011 |
| | ✓ | **0.5642 ± 0.013** | **0.8436 ± 0.012** | **0.6925 ± 0.006** | **0.8456 ± 0.015** | **0.4105 ± 0.014** | **0.6558 ± 0.006** |
| Molformer | × | 0.1238 ± 0.011 | 0.3921 ± 0.004 | 0.1969 ± 0.004 | 0.3901 ± 0.012 | 0.0841 ± 0.010 | 0.2696 ± 0.005 |
| | ✓ | **0.2424 ± 0.015** | **0.6516 ± 0.009** | **0.3850 ± 0.011** | **0.6210 ± 0.014** | **0.1681 ± 0.012** | **0.4579 ± 0.007** |

Apart from that, we provide a full comparison with all existing approaches in Table 2. We elect RWplus [99], ProQ3D [88], VoroMQA [64], SBROD [47], 3DCNN [86], 3DGNN [86], 3DOCNN [65], DimeNet [51], GraphQA [24], and GBPNet [6] as the baselines. Performance is recorded in Table 2, where the second best is underlined. It can be concluded that even if GVP-GNN is not the best architecture, it can largely outperform existing methods including the state-of-the-art no-pretraining method set by Aykent and Xia [6] (*i*.*e*., GBPNet) and the state-of-the-art pretraining results set by Jing et al. [45] and set the new state-of-the-art if it is enhanced by the protein language model.

**Table 2:**
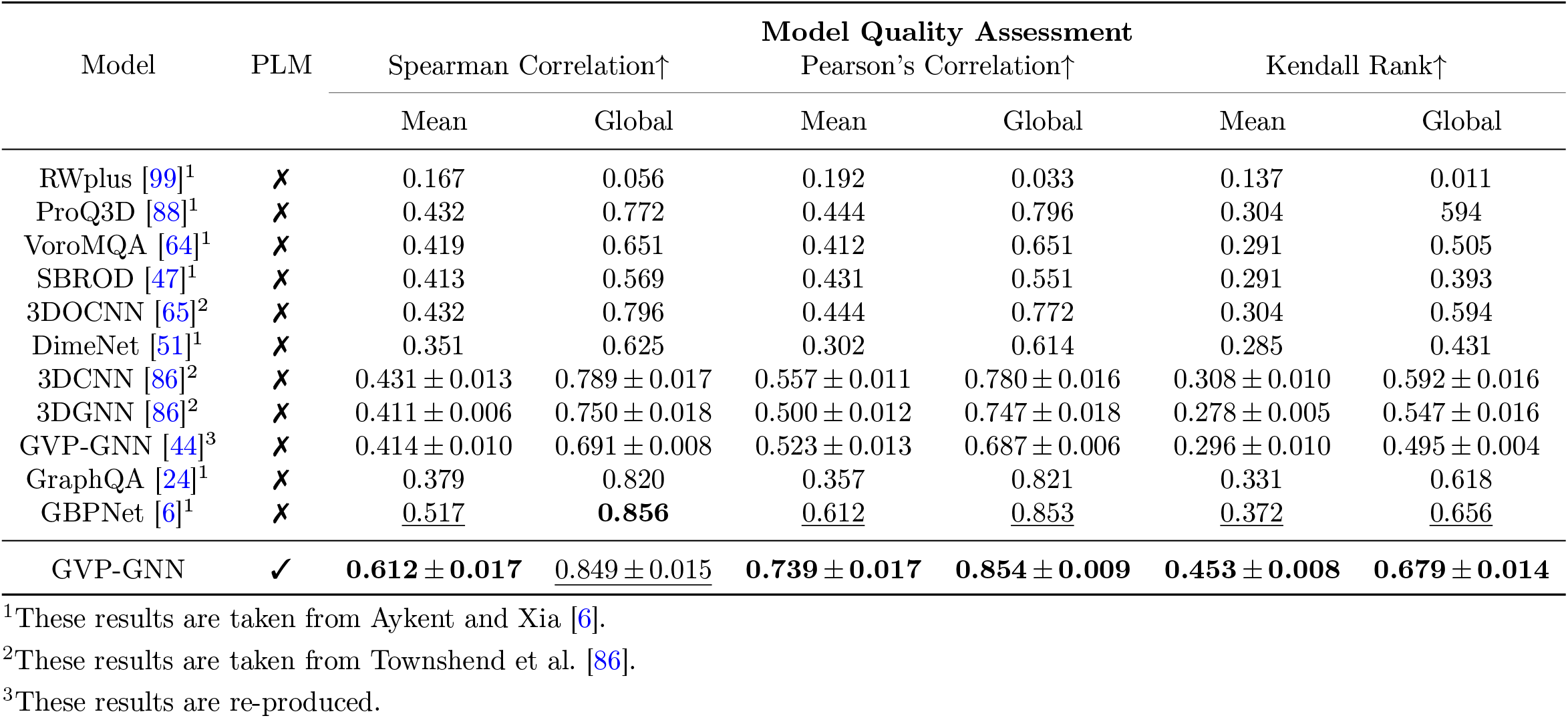
Comparison of performance on MQA. Models are sorted by the year they are released.

#### 2.3.2 Protein-protein Representation Tasks

For PPRD, we report three items as the measurements: the complex root mean squared deviation (RMSD), the ligand RMSD, and the interface RMSD (see Table 3). The interface is determined with a distance threshold less than 8Å. It is noteworthy that, unlike the EquiDock paper, we do not apply the Kabsch algorithm to superimpose the receptor and the ligand. Contrastingly, the receptor protein is fixed during evaluation. All three metrics decrease considerably with improvements of 11.61%, 12.83%, and 31.01% in complex, ligand, and interface median RMSD, respectively. Notably, we also report the result of EquiDock, which is first pretrained on DIPS and then fine-tuned on DB5. It can be discovered that DIPS-pretrained EquiDock still performs worse than EquiDock equipped with pretrained language models. This strongly demon-strates that structural pretraining for GGNNs may not benefit GGNNs more than pretrained protein language models.

**Table 3:** Performance of PPRD on DB5.5 Test Set. Models with ♣ are directly trained and tested on DB5, while EquiDock with ♠ is first pretrained on DIPS and then fine-tuned on the DB5 training set.

| Model | PLM | Protein-protein Rigid-body Docking |  |  |  |  |  |  |  |  |
| --- | --- | --- | --- | --- | --- | --- | --- | --- | --- | --- |
|  |  | Complex RMSD ↓ |  |  | Ligand RMSD ↓ |  |  | Interface RMSD ↓ |  |  |
|  |  | Median | Mean | Std | Median | Mean | Std | Median | Mean | Std |
| EquiDock | ✗♣ | 16.88 | 17.11 | 5.33 | 40.35 | 37.97 | **12.94** | 16.19 | 37.97 | 4.47 |
|  | ✗♠ | 15.02 | 14.31 | 5.28 | 36.82 | 35.95 | 13.18 | 14.37 | 35.68 | **4.12** |
|  | ✓♣ | **14.92** | **13.14** | **4.59** | **35.17** | **33.48** | 14.34 | **11.17** | **33.48** | 4.38 |

For PPI, we record AUROC as the metric in Figure 2. It can be found that AUROC increases for 6.93%, 14.01%, and 22.62% for GVP-GNN, EGNN, and Molformer respectively. It is worth noting that Molformer falls behind EGNN and GVP-GNN originally in this task. But after injecting knowledge learned by protein language models, Molformer achieves competitive or even better performance than EGNN or GVP-GNN. This indicates that protein language models can realize the potential of GGNNs to the full extent and greatly narrow the gap between different geometric deep learning architectures. The results mentioned above are amazing because, unlike MQA, PPRD and PPI study the geometric interactions between two proteins. Though existing protein language models are all trained on single protein sequences, our experiments show that the evolution information hidden in unpaired sequences can also be valuable to analyze the multi-protein environment.

**Fig. 2:**
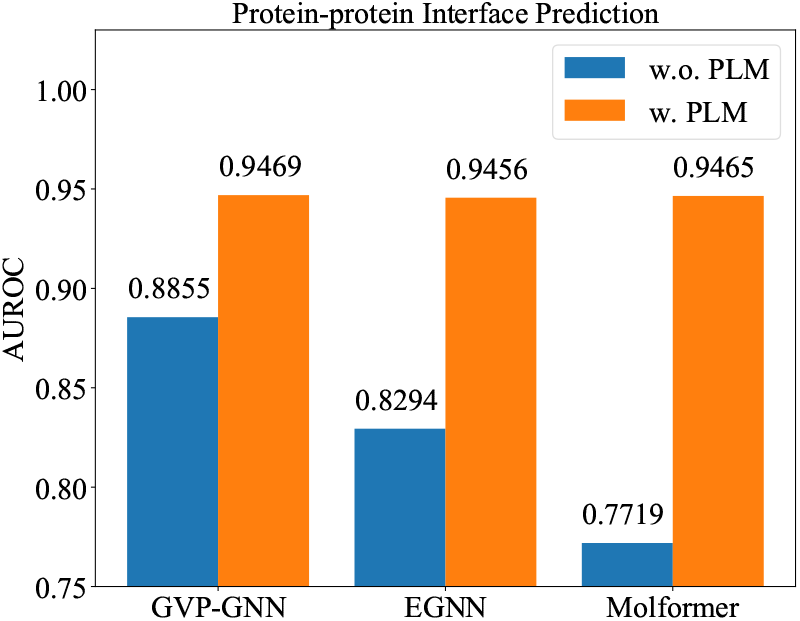
Results of PPI.

#### 2.3.3 Protein-molecules Representation Task

For LBA, we compare RMSD, *R*_*S*_, *R*_*P*_, and *K*_*R*_ in Table 4. The incorporation of protein language models produces a remarkably average decline of 11.26% and 6.15% in RMSD for 30% and 60% identity, an average increase of 51.09% and 9.52% in *R*_*P*_ for the 30% and 60% identity, an average increment of 66.60% and 8.90% in *R*_*S*_ for the 30% and 60% identity, and an average increment of 68.52% and 6.70% in *K*_*R*_ for the 30% and 60% identity. It can be seen that the improvements in the 30% sequence identity is higher than that in the less restrictive 60% sequence identity. This confirms that protein language models benefit GGNNs more when the unseen samples belong to different protein domains. Moreover, contrasting PPRD or PPI, LBA studies how proteins interact with small molecules. Our outcome demonstrates that rich protein representations encoded by protein language models can also contribute to the analysis of protein’s reaction to other non-protein drug-like molecules.

**Table 4:** Results on LBA.

| Model | PLM | Ligand Binding Affinity<br>Sequence Identity (30%) |  |  |  |
| --- | --- | --- | --- | --- | --- |
|  |  | RMSD↓ | Pearson's Correlation↑ | Spearman Correlation↑ | Kendall Rank↑ |
| GVP-GNN | ✗ | 1.6480 ± 0.014 | 0.2138 ± 0.013 | 0.1648 ± 0.009 | 0.1107 ± 0.012 |
|  | ✓ | **1.4556 ± 0.011** | **0.5373 ± 0.010** | **0.5078 ± 0.005** | **0.3495 ± 0.009** |
| EGNN | ✗ | 1.4929 ± 0.012 | 0.4891 ± 0.017 | 0.4725 ± 0.008 | 0.3291 ± 0.014 |
|  | ✓ | **1.4033 ± 0.013** | **0.5655 ± 0.016** | **0.5448 ± 0.005** | **0.3790 ± 0.007** |
| Molformer | ✗ | 1.9107 ± 0.018 | 0.4618 ± 0.014 | 0.4104 ± 0.011 | 0.2812 ± 0.019 |
|  | ✓ | **1.6028 ± 0.020** | **0.5351 ± 0.017** | **0.5372 ± 0.015** | **0.3758 ± 0.016** |
| Sequence Identity (60%) |  |  |  |  |  |
| GVP-GNN | ✗ | 1.5438 ± 0.015 | 0.6608 ± 0.012 | 0.6668 ± 0.0010 | 0.4797 ± 0.014 |
|  | ✓ | **1.5137 ± 0.019** | **0.6680 ± 0.010** | **0.6716 ± 0.008** | **0.4786 ± 0.012** |
| EGNN | ✗ | 1.5928 ± 0.020 | 0.6274 ± 0.013 | 0.6271 ± 0.017 | 0.4498 ± 0.014 |
|  | ✓ | **1.5595 ± 0.022** | **0.6445 ± 0.015** | **0.6463 ± 0.019** | **0.4656 ± 0.019** |
| Molformer | ✗ | 1.8610 ± 0.018 | 0.5528 ± 0.016 | 0.5309 ± 0.015 | 0.3738 ± 0.017 |
|  | ✓ | **1.5926 ± 0.024** | **0.6524 ± 0.018** | **0.6528 ± 0.016** | **0.4367 ± 0.011** |

In addition, we compare thoroughly with all existing approaches for LBA in Table 5. We select a broad range of models including DeepAffinity [48], Cormorant [4], LSTM [11], TAPE [68], ProtTrans [25], 3DCNN [86], GNN [86], MaSIF [31], DGAT [63], DGIN [63], DGAT-GCN [63], HoloProt [79], and GBPNet [6] as the baseline. We report the comparison in Table 5, where the second best is underlined. It is clear that even if EGNN is a median-level architecture, it can achieve the best RMSD and the best Pearson’s correlation when enhanced by protein language models, beating a group of strong baselines including HoloProt [79] and GBPNet [6].

**Table 5:** Comparison of performance on LBA with 30% sequence identity. Models are sorted by the year they are released.

| Model | PLM | Ligand Binding Affinity<br>Sequence Identity (30%) |  |  |  |
| --- | --- | --- | --- | --- | --- |
|  |  | RMSD↓ | Pearson's Correlation↑ | Spearman Correlation↑ | Kendall Rank↑ |
| DeepAffinity [48]<sup>1</sup> | ✗ | 1.893 ± 0.650 | 0.415 | 0.426 | — |
| Cormorant [4]<sup>2</sup> | ✗ | 1.568 ± 0.012 | 0.389 | 0.408 | — |
| LSTM [11]<sup>3</sup> | ✗ | 1.985 ± 0.006 | 0.165 ± 0.006 | 0.152 ± 0.024 | — |
| TAPE [68]<sup>3</sup> | ✗ | 1.890 ± 0.035 | 0.338 ± 0.044 | 0.286 ± 0.124 | — |
| ProtTrans [25]<sup>3</sup> | ✗ | 1.544 ± 0.015 | 0.438 ± 0.053 | 0.434 ± 0.058 | — |
| 3DCNN [86]<sup>1</sup> | ✗ | 1.414 ± 0.021 | 0.550 | 0.553 | — |
| GNN [86]<sup>1</sup> | ✗ | 1.570 ± 0.025 | 0.545 | 0.533 | — |
| MaSIF [31]<sup>3</sup> | ✗ | 1.484 ± 0.018 | 0.467 ± 0.020 | 0.455 ± 0.014 | — |
| DGAT [63]<sup>2</sup> | ✗ | 1.719 ± 0.047 | 0.464 | 0.472 | — |
| DGIN [63]<sup>2</sup> | ✗ | 1.765 ± 0.076 | 0.426 | 0.432 | — |
| DGAT-GCN [63]<sup>2</sup> | ✗ | 1.550 ± 0.017 | 0.498 | 0.496 | — |
| GVP-GNN [44]<sup>4</sup> | ✗ | 1.648 ± 0.014 | 0.213 ± 0.013 | 0.164 ± 0.009 | 0.110 ± 0.012 |
| EGNN [73]<sup>4</sup> | ✗ | 1.492 ± 0.012 | 0.489 ± 0.017 | 0.472 ± 0.008 | 0.329 ± 0.014 |
| HoloProt [79]<sup>3</sup> | ✗ | 1.464 ± 0.006 | 0.509 ± 0.002 | 0.500 ± 0.005 | — |
| GBPNet [6]<sup>2</sup> | ✗ | 1.405 ± 0.009 | 0.561 | **0.557** | — |
| EGNN | ✓ | **1.403 ± 0.013** | **0.565 ± 0.016** | 0.544 ± 0.005 | **0.379 ± 0.007** |

<sup>1</sup>These results are taken from Townshend et al. [86].<sup>2</sup>These results are taken from Aykent and Xia [6].<sup>3</sup>These results are copied from Sonnath et al. [79].<sup>4</sup>These results are re-produced.

### 2.4 Scale of Protein Language Models

It has been observed that as the size of the language model increases, there are consistent improvements in tasks like structure prediction [54]. Here we conduct an additional ablation study to investigate the effect of protein language models’ sizes on GGNNs. Specifically, we explore different ESM-2 with the parameter numbers of 8M, 35M, 150M, 650M, and 3B. Results are plotted in Figure 3, which verifies that scaling the protein language model is advantageous for GGNNs.

**Fig. 3:**
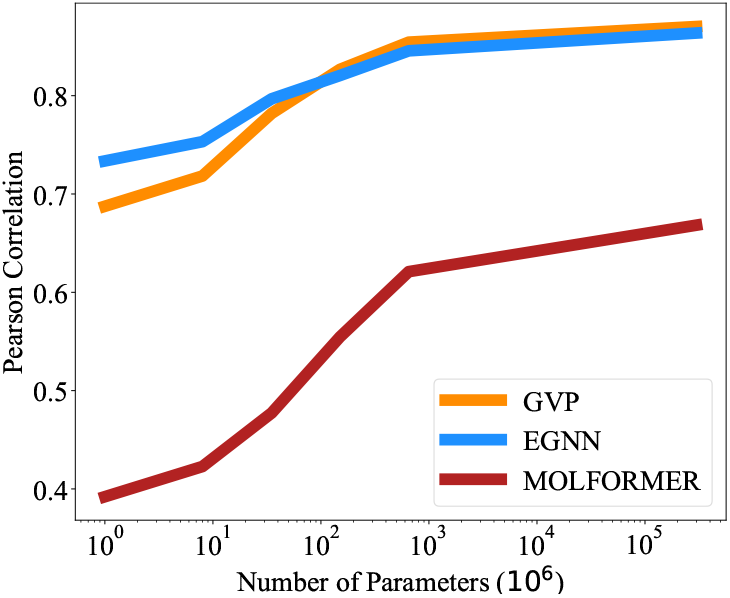
Performance of GGNNs on MQA with ESM-2 at different scales.

## 3 Conclusion

In this study, we investigate a problem that has been long ignored by existing geometric deep learning methods for proteins. That is, how to employ the abundant protein sequence data for 3D geometric representation learning. To answer this question, we propose to leverage the knowledge learned by existing advanced pre-trained protein language models, and use their amino acid representations as the initial features. We conduct a variety of experiments such as protein-protein docking and model quality assessment to demonstrate the efficacy of our approach. Our work provides a simple but effective mechanism to bridge the gap between 1D sequential models and 3D geometric neural networks, and hope to throw light on how to combine information encoded in different protein modalities.

## 4 Method

As discussed before, learning on 3D structures cannot benefit from these large amounts of sequential data. Due to this fact, model sizes of those GGNNs are therefore limited or overfitting may occur [39]. On the contrary, it can be seen, comparing the number of protein sequences in the UniProt database [19] to the number of known structures in the PDB, over 1700 times more sequences than structures. More importantly, the availability of new protein sequence data continues to far outpace the availability of experimental protein structure data, only increasing the need for accurate protein modeling tools.

Therefore, it is straightforward to assist GGNNs with pretrained protein language models. To this end, we feed amino acid sequences into those protein language models, where the state-of-the-art ESM-2 [54] is adopted in our case, and extract the per-residue representations, denoted as 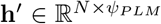. HereΨ_*PLM*_ = 1280. Then **h**^*′*^ can be added or concatenated to the per-atom feature **h**. For residue-level graphs, **h**^*′*^ immediately replaces the original **h** as the input node features.

Notably, incompatibility exists between the experimental structure and its original amino acid sequence. That is, structures stored in the PDB files are usually incomplete and some strings of residues are missing due to inevitable realistic issues [22]. They, therefore, do not perfectly match the corresponding sequences (*i*.*e*., FASTA sequence). There are two choices to address this mismatch. On the one hand, we can simply use the fragmentary sequence as the substitute for the integral amino acid sequence and forward it into the protein language models. On the other hand, we can leverage a dynamic programming algorithm provided by Biopython [18] to implement pairwise sequence alignment and abandon residues that do not exist in the PDB structures. It is empirically discovered that no big difference exists between them, so we adopt the former processing mechanism for simplicity.

## 5 Sequence Recovery Analysis

It is commonly acknowledged that protein structures maintain much more information than their corresponding amino acid sequences. And for decades long, it has been an open challenge for computational biologists to predict protein structure from its amino acid sequence [49, 76]. Though the advancement of Alphafold (AF) [46], as well as RosettaFold [7], have made a huge step in alleviating the limitation brought by the number of available experimentally determined protein structures [40], neither AF nor its successors such as Alphafold-Multimer [26], IgFold [72], and HelixFold [92] are a panacea. Their predicted structures can be severely inaccurate when the protein is orphan and lacks multiple sequence alignment (MSA) as the template. As a consequence, it is hard to conclude that protein sequences can be perfectly transformed to the structure modality by current tools and be used as extra training resources for GGNNs.

Moreover, we argue that even if conformation is a higher-dimensional representation, the prevailing learning paradigm may forbid GGNNs from capturing the knowledge that is uniquely preserved in protein sequences. Recall that GGNNs are mainly diverse in their patterns to employ 3D geometries, the input features include distance [74], angles [50, 51], torsion, and terms of other orders [56]. The position index hidden in protein sequences, however, is usually neglected when constructing 3D graphs for GGNNs. Therefore, in this section, we design a toy trial to examine whether GGNNs can succeed in recovering this kind of positional information.

### 5.1 Protein Graph Construction

Here the structure of a protein can be represented as an atom-level or residue-level graph 𝒢= (𝒱, ℰ), where 𝒢 and ℰ= (*e*_*ij*_) correspond to the set of *N* nodes and *M* edges respectively. Nodes have their 3D coordinates as **x** ∈ ℝ*N ×*3 as well as the initialΨ_*h*_-dimension roto-translational invariant features 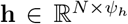 (*e*.*g*., atom types and electronegativity, residue classes). Normally, there are three types of options to construct connectivity for molecules: *r-ball* graphs, *fully-connected* (FC) graphs, and *K-nearest neighbors* (KNN) graphs. In our setting, nodes are linked to *K* = 10 nearest neighbors for KNN graphs, and edges include all atom pairs within a distance cutoff of 8Åfor r-ball graphs.

### 5.2 Recovery from Graphs to Sequences

Since most prior studies choose to establish 3D protein graphs based on purely geometric information and ignore their sequential identities, it provokes the following position identity question:

> *Can existing GGNNs identify the sequential position order only from geometric structures of proteins?*

To answer this question, we formulate two categories of toy tasks (see Figure 4). The first one is a classification task, where models are asked to directly predict the position index ranging from 1 to *N*, the residue number of each protein. This task adopts accuracy as the metric and expects models to discriminate the absolute position of the amino acid within the whole protein sequence.

**Fig. 4:**
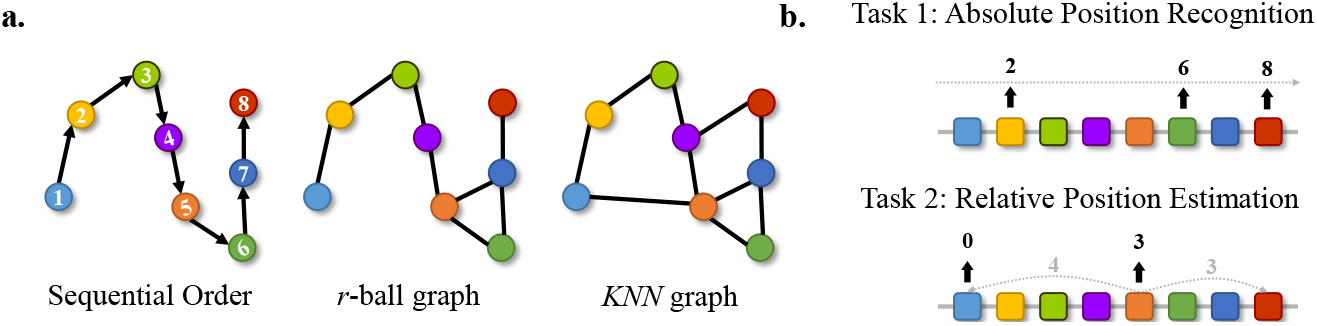
(a) Protein residue graph construction. Here we draw graphs in 2D for better visualization but study 3D graphs for GGNNs. (b) Two sequence recovery tasks. The first requires GGNNs to predict the absolute position index for each residue in the protein sequence. The second aims to forecast the minimum distance of each amino acid to the two sides of the protein sequence.

In addition to that, we propose the second task to focus on the relative position of each residue, where models are required to predict the minimum distance of residue to the two sides of the given protein. We use the root mean squared error (RMSE) as the metric. This task aims to examine the capability of GGNNs to distinguish which segment the amino acid belongs to (*i*.*e*., the center section of the protein or the end of the protein).

### 5.3 Experimental Setting

We adopt three technically distinct and broadly accepted architectures of GGNNs for empirical verification. To be specific, **GVP-GNN** [44, 45] extends standard dense layers to operate on collections of Euclidean vectors, performing both geometric and relational reasoning on efficient representations of macromolecules. **EGNN** [73] is a translation, rotation, reflection, and permutation equivariant GNN without expensive spherical harmonics. **Molformer** [95] employs the self-attention mechanism for 3D point clouds while guarantees SE(3)-equivariance.

We exploit a small non-redundant subset of high-resolution structures from the PDB. To be specific, we use only X-ray structures with resolution *<* 3.0Å, and enforce a 60% sequence identity threshold. This results in a total of 2643, 330, and 330 PDB structures for the train, validation, and test sets, respectively. Experimental details, the summary of the database, and the description of these GGNNs are elaborated in Appendix A.

### 5.4 Results and Analysis

Table 6 documents the overall results, where metrics are labeled with ↑/↓ if higher/lower is better, respectively. It can be found that all GGNNs fail to recognize either the absolute or the relative positional information encoded in the protein sequences with an accuracy lower than 1% and a high RMSE.

**Table 6:** Results of two residue position identification tasks.

| Target | Metric | GVP |  | EGNN |  | Molformer |
| --- | --- | --- | --- | --- | --- | --- |
|  |  | r-ball graph | KNN graph | r-ball graph | KNN graph | FC graph |
| Index | Accuracy (%) ↑ | 0.157 | 0.158 | 0.150 | 0.131 | 0.148 |
| Distance | RMSE ↓ | 392.38 | 392.38 | 412.70 | 403.86 | 270.69 |

This phenomenon stems from the conventional ways to build graph connectivity, which usually excludes sequential information. To be specific, unlike common applications of GNNs such as citation networks [75, 83], social networks [27, 36], knowledge graphs [15], molecules do not have explicitly defined edges or adjacency. On the one hand, r-ball graphs utilize a cut-off distance, which is usually set as a hyperparameter, to determine the particle connections. But it is hard to guarantee a cut-off to properly include all crucial node interactions for complicated and large molecules. On the other hand, FC graphs that consider all pairwise distances will cause severe redundancies, dramatically increasing the computational complexity especially when proteins consist of thousands of residues. Besides, GGNNs also easily get confused by excessive noise, leading to unsatisfactory performance. As a remedy, KNN becomes a more popular choice to establish graph connectivity for proteins [29, 32, 80]. However, all of them take no account of the sequential information and require GGNNs to learn this original sequential order during training.

The lack of sequential information can yield several problems. To begin with, residues are unaware of their relative positions in the proteins. For instance, two residues can be close in the 3D space but distant in the sequence, which can mislead models to find the correct backbone chain. Secondly, according to the characteristics of the MP mechanism, two residues in a protein with the same neighborhood are expected to share similar representations. Nevertheless, the role of those two residues can be significantly separate [62] when they are located at different segments of the protein. Thus, GGNNs may be incapable of differentiating two residues with the same 1-hop local structures. This restriction has already been distinguished by several works [42, 100], but none of them make a strict and thorough investigation. Admittedly, sequential order may only be necessary for certain tasks. But this toy experiment strongly indicates that the knowledge monopolized by amino acid sequences can be lost if GGNNs only learn from protein structures.

## 6 Related Work

### 6.1 Geometric Deep Learning for Proteins

Gigantic molecules (*i*.*e*., macromolecules) populate a cell, providing it with irreplaceable functions for life. And past few years have witnessed growing attention in learning on their 3D structures. Early work adopts graph kernels and support vector machines to classify enzymes based on their conformations [14]. Later, inspired by the booming development of computer vision, protein tertiary structures are represented as 3D density maps with 3DCNNs to address a host of problems such as protein binding site prediction [43], enzyme classification [3], protein-ligand binding affinity [67], protein quality assessment [21], and protein-protein interaction interface identification [85].

At the same time, due to the fact that molecules can be naturally modeled as graphs in real-world studies, GGNNs have emerged as the mainstream line to learn directly from the protein spatial neighboring graphs. They show promising capacity in many tasks including protein interface prediction [29], protein design [42, 45, 81], protein quality assessment [9], function prediction [34], and in more challenging ones like protein folding [39].

Most existing GGNNs rely on the widely adopted message passing (MP) paradigm [20, 33, 56] to aggregate local neighborhood information for the update of node features. Their divergence mainly lies in how to exploit different types of 3D information such as bond lengths or dihedral angles [41, 50, 51, 56]. Moreover, equivariance is regarded as a ubiquitous property for molecular systems, and plenty of evidence has proven the effectiveness to integrate such inductive bias into GGNNs for modeling 3D geometry [4, 10, 30, 84]. Nevertheless, the potential of GGNNs is largely underestimated and cannot be fully released owing to the sparsity of structure data.

### 6.2 Protein Language Modeling

A large body of work has focused on protein language modeling in individual protein families, solving problems like functional nanobody design [78] and protein sequence generation [87]. This success has triggered a prospective trend to model large-scale databases of protein sequences rather than families of related sequences, where unsupervised learning becomes the preferred option. To be explicit, Bepler and Berger [11] combine unsupervised sequence pretraining with structural supervision to generate sequence embeddings. Alley et al. [1] and Heinzinger et al. [38] demonstrate that LSTM language models are able to capture certain biological properties. In the meantime, Rao et al. [68] evaluated a variety of protein language models across a panel of benchmarks concluding that small LSTMs and Transformers fall well short of features from the bioinformatics pipeline. Rives et al. [71] start to model protein sequences with self-attention, illustrating that Transformer-based protein language models can seize accurate information of structure and function in their representations.

A combination of model scale and architecture improvements has been critical to recent successes in protein language modeling. Elnaggar et al. [25] analyze a diversity of Transformer variants. Rives et al. [71] prove that large Transformer models can achieve state-of-the-art features across various tasks. Vig et al. [90] discover that specific attention heads of pretrained Transformers have immediate correlations with protein contacts. Moreover, Rao et al. [69] find that the combination of multiple attention heads is more accurate than Potts models in contact prediction, even if using a single sequence for inference.

For the sake of better capturing the biochemical knowledge, a group of typical pretraining objectives is explored including next amino acid prediction [1, 25], masked language modeling (MLM) [37], contrastive predictive coding [58], and conditional generation [59]. Besides that, Sturmfels et al. [82] and Sercu et al. [77] study alternative objectives with sets of sequences for supervision. Sturmfels et al. [82] extend the unsupervised language modeling to forecast the position-specific scoring matrix (PSSM). Apart from approaches that merely learn in the whole sequence space, multiple sequence alignment (MSA)-based methods leverage the sequences within a protein family to seize the conserved and variable regions of homologous sequences [13, 60, 70].

## Data Availability

The data of model quality assessment, protein-protein interface prediction, and ligand affinity prediction is available by https://www.atom3d.ai/. The data of protein-protein rigid-body docking can be downloaded directly from the official repository of Equidock https://github.com/octavian-ganea/equidock_public.

## Code availability

The code repository is stored at https://github.com/smiles724/bottleneck.

## Authors’ contributions

F.W. and J.X. led the research. F.W. contributed technical ideas. F.W. and Y.T. developed the proposed method. F.W., D.R., and Y.T. performed the analysis. J.X. and D.R. provided evaluation and suggestions. All authors contributed to the manuscript.

## Acknowledgments

This work is supported in part by the Institute of AI Industry Research at Tsinghua University and the Molecule Mind.

## Appendix A Experimental Details

### A.1 Dataset Description

#### Sequence position identification

We use a subset of 3243 high-resolution structures from the PDB and adopt a random split of 2643/330/330 for train/val/test. The distribution of the number of residues is plotted in Figure A1. For the train set, the maximum and the minimum number of residues are 6248 and 43 separately. The mean and standard deviation of the number of residues are 389.9 and 385.7. For the validation set, the maximum and the minimum number of residues are 9999 and 59 separately. The mean and standard deviation of the number of residues are 454.3 and 772.0. For the test set, the maximum and the minimum number of residues are 2326 and 59 separately. The mean and standard deviation of the number of residues are 383.2 and 286.2.

**Fig. A1.**
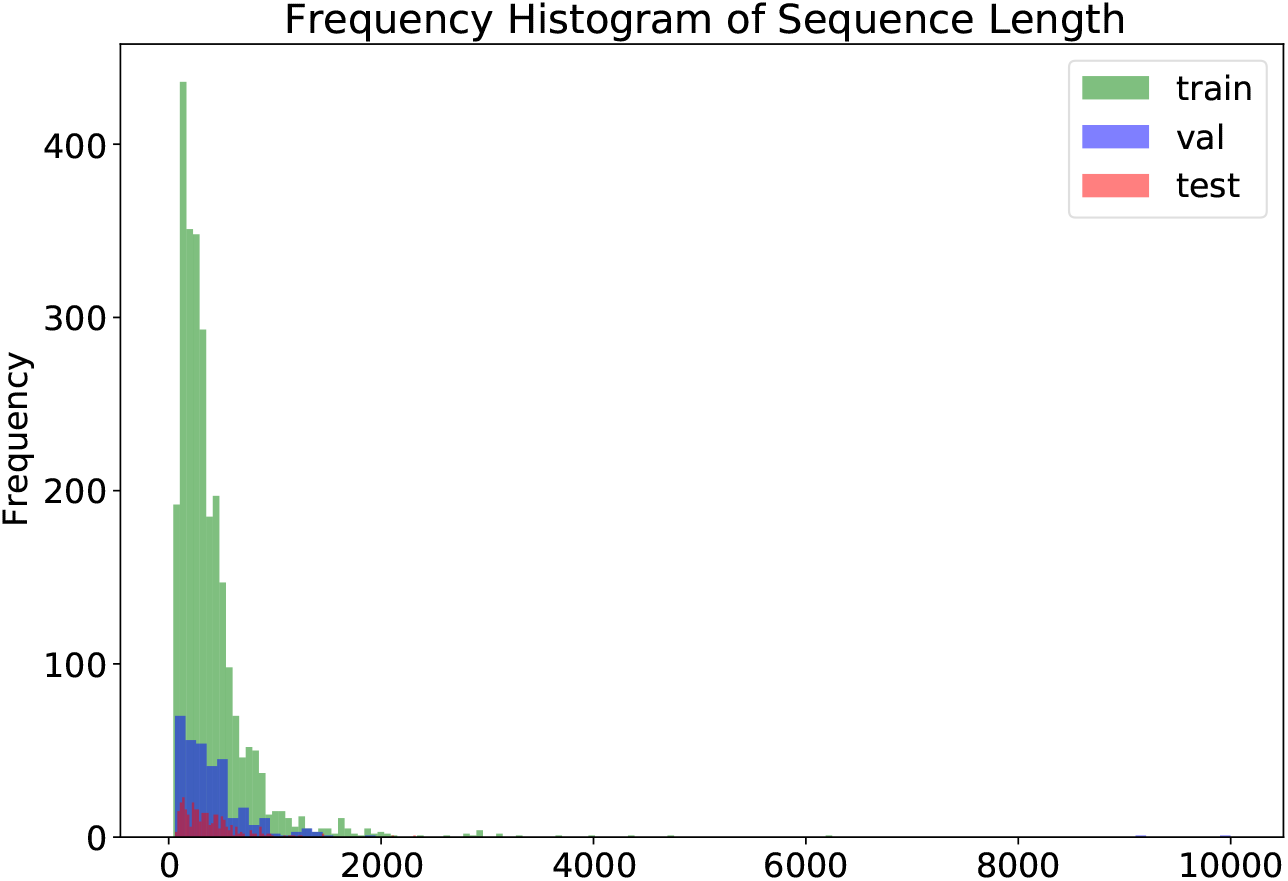
The histogram of the sequence length

#### Model quality assessment

The Critical Assessment of Structure Prediction (CASP) [52] is a long-running international competition held biennially, of which CASP13 is the most recent that addresses the protein structure prediction problem by withholding newly solved experimental structures. Mirroring the setup of the competition, we follow Townshend et al. [86] and split the decoy sets based on target and released year. We choose CASP11 as test set, as the targets in CASP12-13 are not fully released yet. This leads to a dataset split of 25400/2800/16014 for train/val/test.

#### Protein-protein rigid-body docking

We use the DB5.5 databases, which is obtained from https://zlab.umassmed.edu/benchmark/. It is randomly partitioned in train/val/test splits of sizes 203/25/25. It is worth mentioning that DB5.5 also includes unbound protein structures, however, which mostly show rigid structures.

#### Protein-protein interface prediction

We adopt a part of the DIPS database and use a data split of 12216/1526/1526 for train/val/test. Each complex is an ensemble, with the bound ligand and bound receptor structures forming 2 distinct sub-units of said ensemble. We then define the neighboring amino acids as those with any *α*-carbon within 8Åof one another. These neighbors are then included as the positive samples, with all other residues as negatives. At prediction time, we attempt to re-predict which possible residues are positive or negative. In other words, we desire to determine whether each residue is located at the binding pocket. AUROC of those predictions are used as the metric to evaluate performance.

#### Ligand affinity prediction

PDBbind contains X-ray structures of proteins bound to small molecules and peptide ligands, We use the dataset mined from PBDbind by Townshend et al. [86], which has two splits based on 30% and 60% sequence identity thresholds, respectively. Splitting using 30% sequence identity results in train/-val/test of 3507/466/490, while splitting using 60% sequence identity results in train/val/test of 3678/460/460.

### A.2 Backbone Architecture

For tasks that only require the predictive model to output a scalar for the protein/complex or each residue including model quality assessment, protein-protein interface prediction, and ligand binding affinity prediction, we select GVP-GNN, EGNN, and Molformer as the backbone architecture. For more complicated tasks that require more complex computational processes such as protein-protein rigid-body docking, we use specific models to address them like Equidock.

#### GVP-GNN

GVP-GNN [44, 45] is an equivariant GNN in which all node and edge embeddings are tuples (**s, V**) of scalar feature and geometric vector features. Message and update functions are parameterized by *geometric vector perceptrons* (GVPs) – modules mapping between tuples (**s, V**) while preserving rotation equivariance. Its computational process is described in Algorithm 1, where **s** and **V** correspond to the node embedding **h** and coordinates **x** separately. :

##### Algorithm 1

GVP-GNN

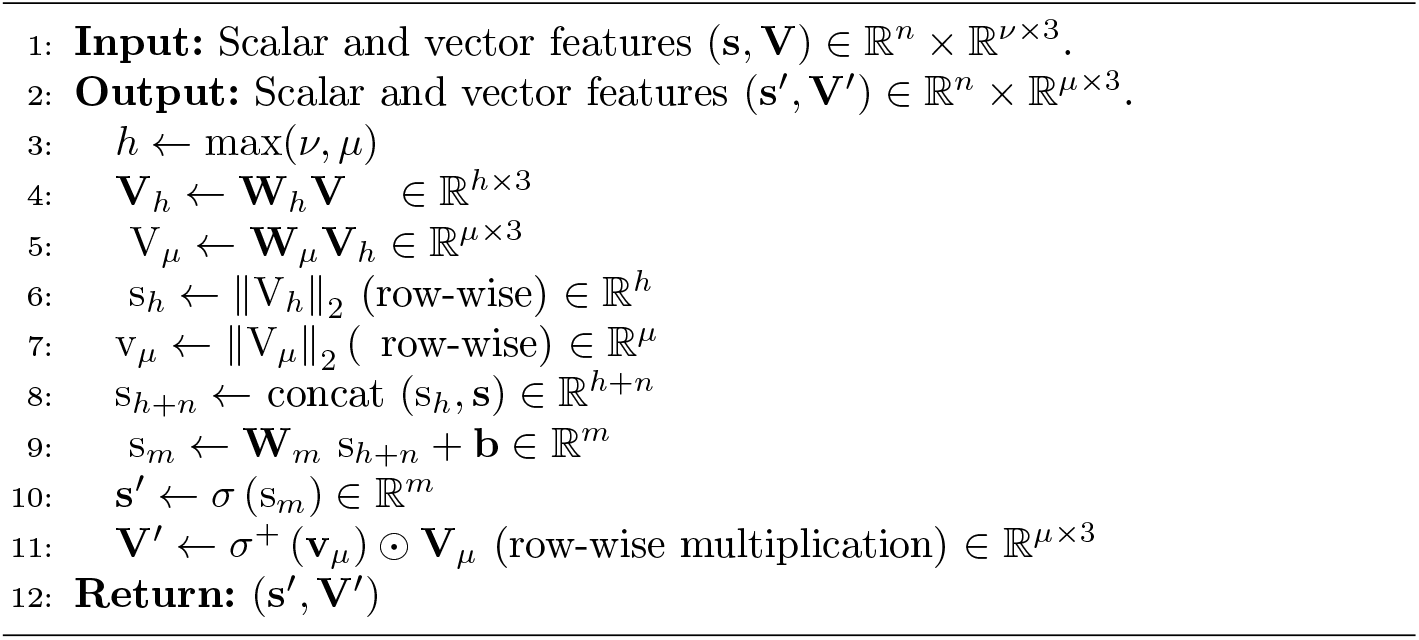

At its core, GVP-GNN consists of two separate linear transformations **W**_*m*_ and **W**_*h*_ for the scalar and vector features, followed by nonlinearities *σ, σ*^+^. An additional linear transformation **W**_*μ*_ is inserted before the vector nonlinearity to control the output dimensionality independently of the number of norms extracted. We adopt a 5-layer GVP-GNN with a dropout rate of 0.7 and a ReLU activation function. The number of radial bases in the edge embedding is 16 and the node dimension is set as (100, 16). All implementation codes are downloaded from the official repository in https://github.com/drorlab/gvp.

#### EGNN

EGNN [73] achieves equivariance without expensive high-order representations in intermediate layers and also realizes competitive performance. Its Equivariant Graph Convolutional Layer (EGCL) takes the set of node embedding 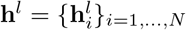, the coordinate embeddings 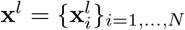 and edge information ℰ = (*e*_*ij*_) as input, and then outputs a transformation on **h**^*l*+1^ and **x**^*l*+1^. Concisely, The equations that define this layer are described as follows:

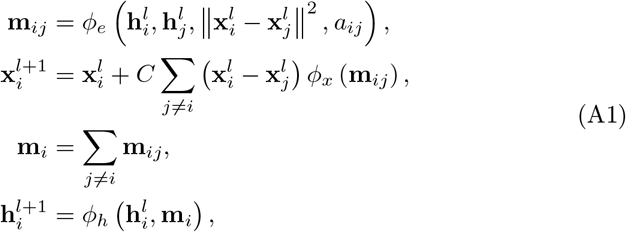

where 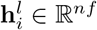 is the nf_dimensional embedding of node *v*_*i*_ at layer *l. a*_*ij*_ are the edge attributes. *ϕ*_*e*_ and *ϕ*_*h*_ are the edge and node operations respectively which are commonly approximated by Multi-layer Perceptrons (MLPs). *ϕ*_*x*_ : ℝ^*nf*^ → ℝ^1^ is the function that takes the edge embedding **m**_*ij*_ as input from the previous edge operation and outputs a scalar value. *C* is chosen to be 1*/* ‖ 𝒩_*I*_ ‖, which divides the sum by its number of neighboring (connected) elements. We choose a 4-layer EGNN with a Siwsh activation function as a non-linearity. The number of the dimension for edges is 16 and residue connections are used. All implementation codes are downloaded from the official repository in https://github.com/vgsatorras/egnn.

#### Molformer

Molformer [95] is a variant of Transformer that employs a heterogeneous self-attention layer to differentiate the interactions between multi-level nodes. Here we use a weaker version of Molformer. To be explicit, we do not extract any sort of motifs from either protein or small molecules. Besides, we abandon the multi-scale self-attention mechanism and the Attentive Farthest Point Sampling (AFPS) for more efficient computations and only use the global features. Even though we pick up a simplified form of Molformer, its performance is competitive with or even outperforms EGNN and GVP-GNN on all tasks. The Molformer architecture has 2 layers, 4 heads, a hidden-layer dimension of 1280, and a dropout rate of 0.1. All implementation codes are downloaded from the official repository in https://github.com/smiles724/molformer.

#### EquiDock

Equidock [32] predicts the rotation and translation to place on of the proteins at the right docked position relative to the second protein. It adopts an Independent E(3)-Equivariant Graph Matching Network (IEGMN), which extends both Graph Matching Networks (GMN) and EGNN. It performs node coordinate and feature embedding updates for an input pair of protein graphs and uses inter- and inter-node messages, as well as E(3)-equivariant coordinate updates. The backbone IEGMN has 5 layers and no dropout. It uses LeakyReLU as the activation function. and does not use distance as an edge feature. All implementation codes are downloaded from the official repository in https://github.com/octavian-ganea/equidock_public.

### A.3 Training Details

We run all experiments on 2 A100 GPUs, each with a memory of 80G. For MQA, PPRD and PPI, we use residue-level graphs for protein representation learning. For PPRD, models are trained using Adam with a learning rate of 2e-4 and early stopping with patience of 30 epochs. For MQA, PPI, and LBA, models are trained using Adam with a learning rate of 1e-4 and early stopping with patience of 8 epochs. A Plateau learning rate scheduler is applied with a factor of 0.6, patience of 5, and a minimum learning rate of 5e-7. The batch size is 32 if no out-of-memory (OOM) error is not triggered, otherwise, we adopt a batch size of 16. The maximum epoch is set as 200.

For PPRD, we randomly assign the roles of ligands and receptors during training. For PPI, as mentioned before, we formulate interface as the residues whose least distances to their counterpart protein are within 8 Å. For LBA, we only use the residues within a distance of 6 Å from the ligand (*i*.*e*., the pocket) following Townshend et al. [86]. We build heterogeneous molecular graphs where nodes for proteins are residue-level and nodes for ligands are atom-level. Here we do not distinguish different atom types and simply regard all atoms as the same group, which is denoted as the new ‘LIG’ pseudo residue class.

For the protein language model, we use the ESM-2 with a parameter size of 650M and 33 layers as the default one. It is trained on UR50/D2021_04 and has an embedding dimension of 1280. We extract per-residue representations as the input for each task. For the ablation study, we adopt ESM-2 with parameter sizes of 8M, 35M, 150M, and 3B that are all trained on the same UR50/D2021_04 dataset with different layers of 6, 12, 30, and 36 respectively. For more details, please visit the official website of ESM in https://github.com/facebookresearch/esm.

## Appendix B Limitations

In spite of our successful confirmation that protein language models can promote geometric deep learning, there are several limitations and extensions of our framework left open for future investigation. First, our work only examines the efficacy of the state-of-the-art protein language model, ESM-2. It should be more convincing if more types of protein language models like MSA-Transformer and ProtTrans are verified. Second, our 3D protein graphs are residue-level. We believe atom-level protein graphs also benefit from our approach, but its increase in performance needs further exploration.

